# Deciphering microbiome impacts on fungal-microalgal interaction in a marine environment using metabolomics

**DOI:** 10.1101/2021.05.27.445989

**Authors:** Olivier Berry, Enora Briand, Alizé Bagot, Maud Chaigne, Laurence Meslet-Cladière, Julien Wang, Olivier Grovel, Jeroen J. Jansen, Nicolas Ruiz, Thibaut Robiou du Pont, Yves François Pouchus, Philipp Hess, Samuel Bertrand

**Affiliations:** Université de Nantes, MMS, EA 2160, Nantes, France; IFREMER, DYNECO, Laboratoire Phycotoxines, F-44000 Nantes, France; Laboratoire Universitaire de Biodiversité et Écologie Microbienne, Université Brest, F-29280 Plouzané, France; Radboud University, Institute for Molecules and Materials, Heyendaalseweg 135, 6525 AJ Nijmegen, The Netherlands

**Author notes:** Corresponding author: Samuel Bertrand, Laboratoire Mer, Molécules, Santé – EA 2160, UFR des Sciences Pharmaceutiques et Biologiques, Université de Nantes, Nantes, France.

**Keywords:** *Aspergillus pseudoglaucus*, metabolomics, microbial co-culture, microbiome, *Prorocentrum lima*

## Abstract

The comprehension of microbial interactions is one of the key challenges in microbial ecology. The present study focuses on studying the chemical interaction between the toxic dinoflagellate *Prorocentrum lima* PL4V strain and associated fungal strains (two *Penicillium* sp. strains and three *Aspergillus* sp) among which the *Aspergillus pseudoglaucus* strain MMS1589 was selected for further co-culture experiment. Such rarely studied interaction (fungal-microalgal) was explored in axenic and non-axenic conditions, in a dedicated microscale marine environment (hybrid solid/liquid conditions), to delineate specialized metabolome alteration in relation to the *P. lima* and *A. pseudoglaucus* co-culture in regard to the presence of their associated bacteria. Such alteration was monitored by high-performance liquid chromatography coupled to high-resolution mass spectrometry (LC-HRMS). In-depth analysis of the resulting data highlighted (1) the chemical modification associated to fungal-microalgal co-culture, and (2) the impact of associated bacteria in microalgal resilience to fungal interaction. Even if only a very low number of highlighted metabolites were fully characterised due to the poor chemical investigation of the studied species, a clear co-culture induction of the dinoflagellate toxins okadaic acid and dinophysistoxin 1 was observed. Such results highlight the importance to consider microalgal microbiome to study parameters regulating toxin production. Finally, a microscopic observation showed an unusual physical interaction between the fungal mycelium and the dinoflagellates.

## Introduction

In the marine environment, microalgae are in constant interaction, both at the intra- and inter-specific levels, which span from competition, mutualism, symbiosis, parasitism and predation [1]. These large diversity of biotic interactions are governed by trophic relationships (*e*.*g*. nutrients acquisition) and chemical communication mediated by diffusible signalling metabolites [2, 3]. Among those marine microalgae, toxic dinoflagellates represent particular microorganisms that may drastically impact marine ecosystems, *e*.*g*. reduction of fishes and algae diversity, as well as, human health hazards (seafood contamination) [4, 5]. Harmful algal blooms (HABs) are considered detrimental events, and HAB species are increasingly observed in some ecosystems, especially at human activities’ proximity [4]. One HABs-related threat is diarrheic shellfish poisoning which is caused by toxins, like okadaic acid (OA). These latter polyketide toxins are produced by various marine dinoflagellates, including *Dinophysis* spp. and *Prorocentrum* spp. Such production is known to be influenced by nitrogen levels, light intensity, temperature and/or salinity [6,7]. The high energy investment required by the biosynthesis of such high molecular weight molecules point out the possible fitness advantage these toxins procure to their producers [8].

*Prorocentrum lima* has benthic lifestyle [9], which implies complex mechanisms to compete for nutrients, light and space [10]. Such competition requires the production of specialized metabolites, which are often evocated or revealed indirectly, while only a few studies highlight such molecules. This is partly due to technical difficulties appearing in their isolation and subsequent characterisation (*e*.*g*. quantities). Besides abiotic inducers, OA is involved in the toxic response to *Artemia salina* chemical cues. The exposition of *P. lima* to such predator yields OA over-production, and following impaired survival rates in predator population [11]. However, it remains unclear whether OA exerts a broad spectrum deterrence effect, since only a few number of studies directly linked OA exposure to negative predator outcomes [12]. As an example, such response is also observed with *Alexandrium minutum* in response to the copepod *Acartia tonsa* which produces copepodamides (polar lipids) yielding an increase up to 20-fold of parasitic shellfish toxin production [13]. Besides predators, co-occurring dinoflagellates are also competitors [14]. For instance, *P. lima* was shown to produce a complex chemical mixture (including OA) possessing growth inhibitory against other co-occurring species [15]. Therefore, *P. lima* seems equipped with various competition chemical signals suited to its benthic epiphytic lifestyle.

Aside from predators and competitors, microalgae are known to interact with various bacteria with mutualistic or defensive behaviours. They can alter, through specific metabolite production, the composition of co-occurring bacterial communities [16], or the bacterial metabolism, including quorum sensing [17]. In parallel, bacteria interfere with microalgal physiology, *e*.*g*. through chemical signals from their quorum sensing systems. For example, *Pseudoalteromonas piscicida* releases 2-heptyl-4-quinolone when co-cultured with *Emiliana huxleyi*, causing its higher mortality rates [18]. Moreover, mutualistic evidence in microalgal-bacterial interactions has been shown [19–21]. One example is microalgal dependency on vitamin B12. Some microalgae, living with vitamin B12 bacteria producers, have lost their costly methionine synthase pathway for a less costly vitamin B12 dependent one. Consequently, such microalgae rely entirely on symbiosis with bacteria producing vitamin B12 to be able to grow [19]. In return, bacteria benefit from microalgal carbonated exudates [20]. While there is clear evidence of associated bacteria effect on microalgal metabolism, alteration of toxin production is still unclear [22–24] and only supported by few studies [25, 26].

Meanwhile, in fungi parasitic microscopic chytrid zoospores are known for using microalgae carbohydrates exudates to detect preys [27] and diatoms are able to alter these parasite life cycles after first fungal induced wound, by fast metabolism adaptation towards production of lipid derived polyenals and associated aldehydes [28]. An example of parasite induced toxin production is the up-regulation and production of microalgal self-growth-inhibitory alkaloids when invaded by *Lagenisma coscinodisci* [29]. Interestingly, no data is available on the existence of chemical interactions between saprophytic marine fungi and microalgae while physical associations are already reported to induce microalgal aggregation in bioprocesses [30].

The study of such microbial interaction *in vitro*, remains a very challenging process [31]. Up to now, the best strategies rely on exploring bipartite interaction in controlled conditions [32]. Metabolomics, defined as a non-selective, comprehensive analytical approach for the identification and quantification of metabolites in a biological system [33], remains a method of choice to highlight chemical modification induced by microbial co-culture [33, 34]. Various examples already showed the induction of specific metabolites in microbial co-culture [34] through the expression of hidden biosynthesis gene clusters (BGCs) [35].

This study of chemical mediation between microalgae and saprophytic fungi in a marine environment, focuses on the toxin producer *P. lima* PL4V strain (known to produce OA and dinophysistoxin 1 – DTX-1). It aims at exploring (1) the presence of fungal strains within the microbiome of this epiphytic dinoflagellate, (2) the chemical mediation between *P. lima* and a fungal strain (namely *Aspergillus pseudoglaucus* MMS1589) co-occurring in culture and (3) whether the fungus may influence microalgal toxin production. Therefore, after fungal strain isolation from the *P. lima* PL4V strain, chemical modifications related to microbial co-cultures were highlighted through a metabolomic approach using a high-performance liquid chromatography coupled to high-resolution mass spectrometry (LC-HRMS).

## Materials and Methods

### Microalgae

*Prorocentrum lima* is a benthic dinoflagellate that may be considered a species complex as there is both temperate and tropical strains reported for this species [36]. The *P. lima* PL4V strain was collected by colleagues from the Instituto Espanol Oceanografico (IEO, Vigo) in the lagoon *Lago dos Nenos* at the junction of the two Cíes Islands Illa de Monteagudo and Illa de Faro (Ria de Vigo, Galicia, Spain) [37], a temperate area, and maintained at the French Research Institute for the Exploitation of the Sea (IFREMER).

### Culture media

All media used for fungal strain isolation were prepared using Reef Crystal mixture at 36 g/L (enriched marine salt from Aquarium Systems) to meet standard sea water salt concentrations and autoclaved at 121°C during 20 min. Chloramphenicol (50 mg/mL, Poly Labo, Paul Block & Cie, Strasbourg, France) was added in liquid or solid media used for fungal strain isolation, to avoid bacterial growth. Malt Extract (ME) liquid media was prepared with following concentrations: glucose 20 g/L, peptone (Biokar Diagnostics, Beauvais, France) 1 g/L, Malt Extract (Conda pronadisa, Madrid, Spain) 30 g/L, copper sulphate 5 mg/L, zinc sulphate 1 mg/L. The Dextrose Casein (DC) medium was prepared as follow: glucose 40 g/L, peptone 10 g/L. Solid equivalent of these media were also prepared. Dextrose Casein Agar (DCA) used DCA mixture (Becton, Dickinson and Company, Sparks, NV USA) 65 g/L and the Malt Extract Agar (MEA, Biokar Diagnostics) medium was composed of glucose 20 g/L, peptone 1 g/L, Malt Extract mixture (Conda pronadisa) 45 g/L, copper sulphate 5 mg/L, zinc sulphate 1 mg/L.

For the co-culture experiment, f/2 microalgal growth medium, without silica addition, was used as previously described [38]. The solid f/2 medium (f/2GA) was prepared using f/2 liquid medium complemented by 20 g/L of agar and 2 g/L of glucose. In some cases, large spectrum antibiotics mixture was added to f/2 liquid medium to provide the following final concentration in the culture medium: ampicillin 500 µg/mL, gentamycin 100 µg/mL, kanamycin 200 µg/mL [39].

### Fungal isolation from microalgal biomass

To isolate fungal strains, *P. lima* PL4V biomass was inoculated on several liquid and solid culture media (ME, DC, MEA and DCA media). Fungal growth at 27 °C was evaluated every day. When fungal colonies were observed, each colony was further collected and inoculated on DCA medium for storage and morphological identification based on macroscopic and microscopic features. The MMS1589 strain was further identified based on molecular biology information (ITS and *β*-tubulin sequencing) according to previously published protocol [40]. The ITS and *β*-tubulin sequences for MMS1589 strain were deposited in GenBank under accession numbers MN134000.1 and MN164633.1, respectively.

### Co-culture growth in miniaturised liquid/solid environment

The miniaturised liquid/solid environment was prepared in tilted 50 mL flasks (Figure 1). Firstly, 10 mL f/2GA medium were added in the flask. Subsequently, the *A. pseudoglocus* MMS1589 strain was inoculated by spiking it into the middle of the f/2GA medium. After five days of fungal growth at 24°C under 12h/12h dark-light cycle using artificial light (DENNERLE Nano Light 11 watt, 6000 Kelvin, 900 lumen), 10 mL liquid f/2 medium (with or without antibiotic cocktail), and 15 mL *P. lima* PL4V inoculum in f/2 medium at 30 600 cells/mL were added to the flask. The co-culture was then incubated for six days under 12h/12h dark-light cycle. In parallel, the same design was used for both microorganisms alone (monoculture), where only *A. pseudoglocus* MMS1589 or *P. lima* PL4V was inoculated. In addition, blank conditions were performed without microorganism inoculation.

**Figure 1.**
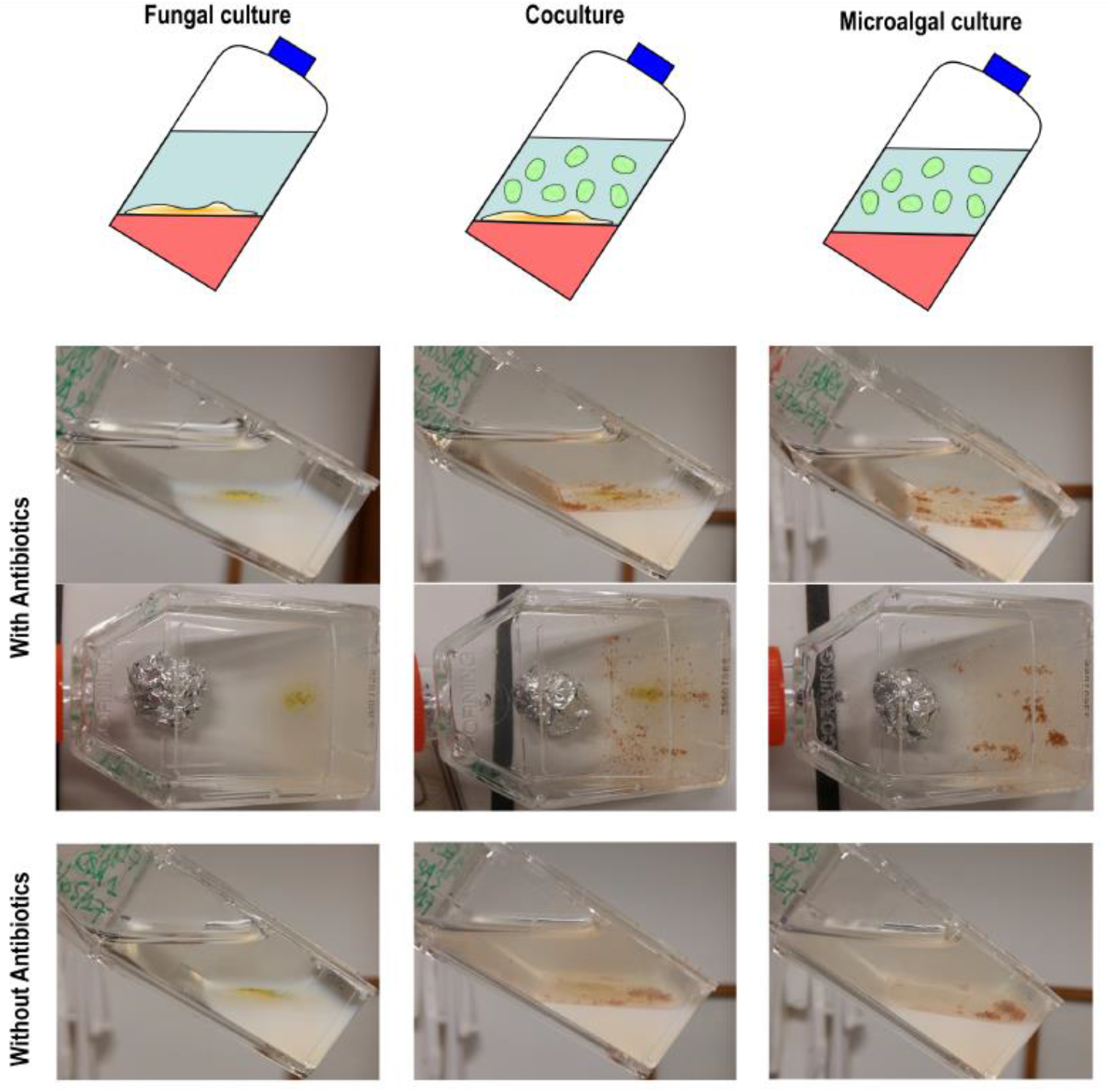
Experimental design of microscale co-culture experiments of marine benthic chemical interactions. Notice the clarity of the water when microalgal inoculum is treated with antibiotics.

Five replicates of each experiment were performed, leading to thirty samples (with antibiotics: five co-cultures, five *A. pseudoglocus* and five *P. lima* monocultures; without antibiotics: five co-cultures, five *A. pseudoglocus* and five *P. lima* monocultures).

### Biomass extraction

After six days, for each growth experiment in liquid/solid environment, the liquid overlaying phase (LOP) was separated from the solid phase (resulting f/2 GA medium and fungus/microalgae) and further centrifuged to remove any cellular debris. Each LOP was extracted three times with 5 mL ethyl acetate (EtOAc), and the three resulting extracts were pooled and dried under vacuum using SpeedVac™ concentrator (Thermo Scientific). The solid phase was extracted twice with 5 mL EtOAc using an ultrasonic bath for 10 min prior to filtration. The two solid phase extracts were pooled and dried under vacuum using a SpeedVac™ concentrator. All extracts were weighed after evaporation to dryness.

### LC-HRMS profiling

Each extract was diluted to a concentration of 1 mg dry weight/mL in HPLC-grade methanol (Biosolve Chimie, Dieuze, France). LC-HRMS profiling was achieved using a UFLC-ESI-IonTrap-TOFMS system (Shimadzu, Marne-la-Vallée, France) according to a previously published protocol [41]. Briefly, chromatographic separation was achieved using a Kinetex™ C_18_ column (100 × 2.1 mm, 2.6 µm), at 40 °C with a 0.3 mL/min flow rate. The mobile phase is a gradient starting at 15 % acetonitrile (B) for 2 min, followed by an increase to 100 % over 23 min and further maintained for 5 min. Mass detection was set between *m/z* 100-1000 with fast switching between positive and negative ionisation mode (PI and NI).

In the sequences, samples were injected randomly using a dedicated Excel macro [42] and MeOH blanks and QC samples were injected every 15 samples. The QC sample was prepared by mixing 30 µL of all samples together.

### Data Analysis

LC-HRMS data were exported as CDF files using LC Solution (Shimadzu) to allow automatic peak picking using MZmine 2 (ref.43). The mass detection was performed using a centroid algorithm with a noise level of 1.5E4. The chromatogram building used a minimum time span of 0.03 min, group intensity threshold of 1E5, a minimal of intensity of 1.5E4 and *m/z* tolerance of 30 ppm [44]. The peak deconvolution was performed using the baseline cut-off algorithm with median *m/z* centre calculation, a peak duration range between 0.05 and 10 min. The chromatograms were deisotoped with *m/z* tolerance of 1 mDa, retention time tolerance of 0.1 min and maximum charge of 3. The peak list was aligned using the Join aligner algorithm with *m/z* tolerance of 1 mDa or 30 ppm, retention time tolerance of 0.2 min, using the same weight for retention time and *m/z* importance. Subsequently, the gap-filling step was performed with a peak-finder algorithm, intensity tolerance of 1.0 *m/z* tolerance of 1 mDa or 30 ppm, retention time tolerance of 0.2 min. The results were exported as a .csv file containing all peaks observed and referenced by their mass to charge ratio (*m/z*) and retention times (t_R_) together with their respective peak areas in each sample, yielding the data matrix.

The peaks detected in blanks were withdrawn from the data matrix as background signal. The absence of ions corresponding to antibiotics and their possible anticipated metabolites using Biotransformer [45] (with phase I, II CYP450 and microbiota transformation) were confirmed. Then, the data matrix containing 6627 features was transformed to reflect full compound production by multiplying each peak area by the corresponding extract amount.

Finally, all data from one culture were concatenated to provide the final data matrix, thus data from replicated sample contained features (*m/z* at t_R_ with corrected peak areas) obtained from LOP extract profiles (NI and PI) and from solid phase extract profiles (NI and PI). Thus, all 30 samples were characterised by 13254 features.

The final data matrix was statistically analysed using R 4.0 (CRAN) with the PocheRmon function [46]. This function allows to analyse co-culture data univariatly using the stats package [47], and multivariatly using both the ropls package [48] and the Projected Orthogonalized CHemical Encounter MONitoring (POChEMon) approach [49] encoded in R. All these approaches allowed to highlight statistically significant features. Highlighted features were confirmed manually in the raw LC-HMS data to remove false peaks detected during the automated peak detection step.

### Annotation of statistically relevant features

The highlighted features were further annotated using traditional dereplication strategy [50] based on mass and spectral accuracy of the MS spectra. Accurate masses (up to 30 ppm for smaller peaks) were searched after adduct mass correction ([M+H]^+^, [M+Na]^+^, [M-H]^-^, [M+Cl]^-^ and [M+formic acid-H]^-^ were considered [51] for compounds reported to be produced by either *Prorocentrum* sp. and *Aspergillus* sp. (also considering its teleomorph *Eurotium* sp. [52]) in the Dictionary of Natural Products® (CRC Press, v.29.2) [53]. Furthermore, OA standard (Diagnostic Chemicals Limited, Charlottetown, CA USA) was injected in the LC-HRMS method and was detected at 14.62 min.

## Results

### Filamentous fungi were isolated from *Prorocentrum lima* fresh biomass

To evaluate the presence of fungi in *P. lima* microbiome, biomass from PL4V strain was used as inoculum for cultivation in fungal adapted medium [54], in presence of antibiotics. Cultivation was achieved either in liquid or solid conditions (Table S1). After few weeks, fungal colonies were observed and isolated (Figure S1). All strains were identified based on fungal morphology resulting in the isolation of two *Penicillium* sp. strains (MMS1593, MMS1596) and three *Aspergillus* sp. strains (MMS1589, MMS1591, MMS1594).

Among isolated strains, the *Aspergillus* sp. strains MMS1589 was selected for further co-culture study, due to its unusual morphology (cleistotheca and ascopores simultaneous observation) (Figure S2) (only one was selected due to expected difficulties in such experimental design, as such fungal-dinoflagellates co-culture were not previously reported). Congo red coloration revealed hyaline septate mycelium and hyaline conidiophores, bearing uniseriate totally covered vesicles. Conidia were round shaped. Interestingly, globose cleistotheca was also visible. Asci and ascopores were present next to cleistotheca. On the basis of the genes encoding fungal small subunit ribosomal RNA (SSU rRNA), Internal Transcribed Spacer 1 (ITS) and β-tubulin of MMS1589, phylogenetic analyses were performed with all sequences in Genebank. The MMS1589 ITS partial gene has 99 % of identity with the *A. pseudoglaucus* strain FJAT-31014 SSU rRNA (partial sequence) and the MMS1589 β-tubulin partial gene has 99 % of identity with the *A. pseudoglaucus* benA gene for β-1 and β-2 tubulin (partial sequence). Finally, this strain was identified as *A. pseudoglaucus*, formerly known as *Eurotium repens* [52].

### Specific microscale marine environment was designed for co-culture

The study of chemical interaction between *P. lima* PL4V and *A. pseudoglaucus* MMS1589, implied the challenging design of a functional micro-environment for microalgal / fungal co-culture since both species have specific requirements for their growth. On one hand, the benthic dinoflagellate *P. lima* growth requires a liquid media (typically f/2 medium) without stirring, without organic carbon source and with a light source for night-and-day cycle. In that case, it grew as a biofilm in the bottom of the culture flask. On the other hand, fungal growth requires either liquid or solid media with an organic carbon source. However, *A. pseudoglaucus* growth in liquid medium without stirring yielded fungal colonies floating over the liquid media. Thus, it was decided to culture the fungus on solid medium (containing a low level of glucose, 2 g/L) prior to addition of liquid medium in the culture flask and *P. lima* inoculation.

When both organisms were cultivated separately in the composite liquid/solid environment with light (12h/12h night-and-day cycle), normal growth was observed (Figure 1): (1) increase of the fungal colony size at the bottom of the flask (on the surface of the agar layer) and (2) formation of a *P. lima* biofilm as usually observed in usual conditions. During *P. lima* culture, bacterial development was observed leading to a cloudy culture medium (Figure 1), as confirmed by microscopic observation (most likely due to the sugar content of the underlying solid medium and the microalgae derived organic matter). Thus, to study microalgal growth with a very limited bacterial development, antibiotics were added in parallel [39].

During co-culture, both microorganisms grew normally (based on morphologic observation). However, *P. lima* tended to gather within the fungal mycelium, possibly using it as anchor to stick to the bottom of the flask. Again, bacterial development was observed in absence of antibiotics in the liquid medium.

### *Aspergillus pseudoglaucus* MMS1589 impacts specialized metabolite production by *Prorocentrum lima* PL4V and its associated bacteria during co-culture

The co-culture experiment was achieved between *P. lima* PL4V and *A. pseudoglaucus* MMS1589 in the devised marine microscale environment in presence and absence of antibiotics [39] to be able to delineate the clear effect *A. pseudoglaucus* on *P. lima*. After 6 days, culture was stopped and agar (bottom part) and liquid supernatant were collected and extracted separately. The agar samples contained fungal mycelium and/or dinoflagellates in a biofilm and the liquid supernatant contained only *P. lima* when inoculated (fungal spores traces were observed). To obtain a clear view of metabolites present in both types of extracts, they were profiled by LC-HRMS [41]. For sake of clarity, single strain *P. lima* culture corresponds to the dinoflagellates with (xenic) or without (axenic) their associated bacteria depending on the presence of antibiotics.

To explore the LC-HRMS data, all peaks were automatically detected using dedicated open-source software [43]. All peaks detected in both LOP and solid phase were concatenated (per samples), generating the final data matrix. The resulting data matrix was subjected to unsupervised statistical analysis through principal component analysis (PCA) to assess the self-organized metabolome differences between *P. lima* PL4V and *A. pseudoglaucus* MMS1589 monocultures, and their corresponding co-culture (Figure 2A). The first two components of the resulting PCA model correspond to 22.1% (PC1) and 11.3% (PC2) of explained variance. This model clearly highlighted the presence of 3 clusters related to *P. lima, A. pseudoglaucus* and their co-culture. The first component (PC1) is related to the differences between the *P. lima* and *A. pseudoglaucus* monocultures, and the second component clearly differentiates between monocultures and co-cultures.

**Figure 2.**
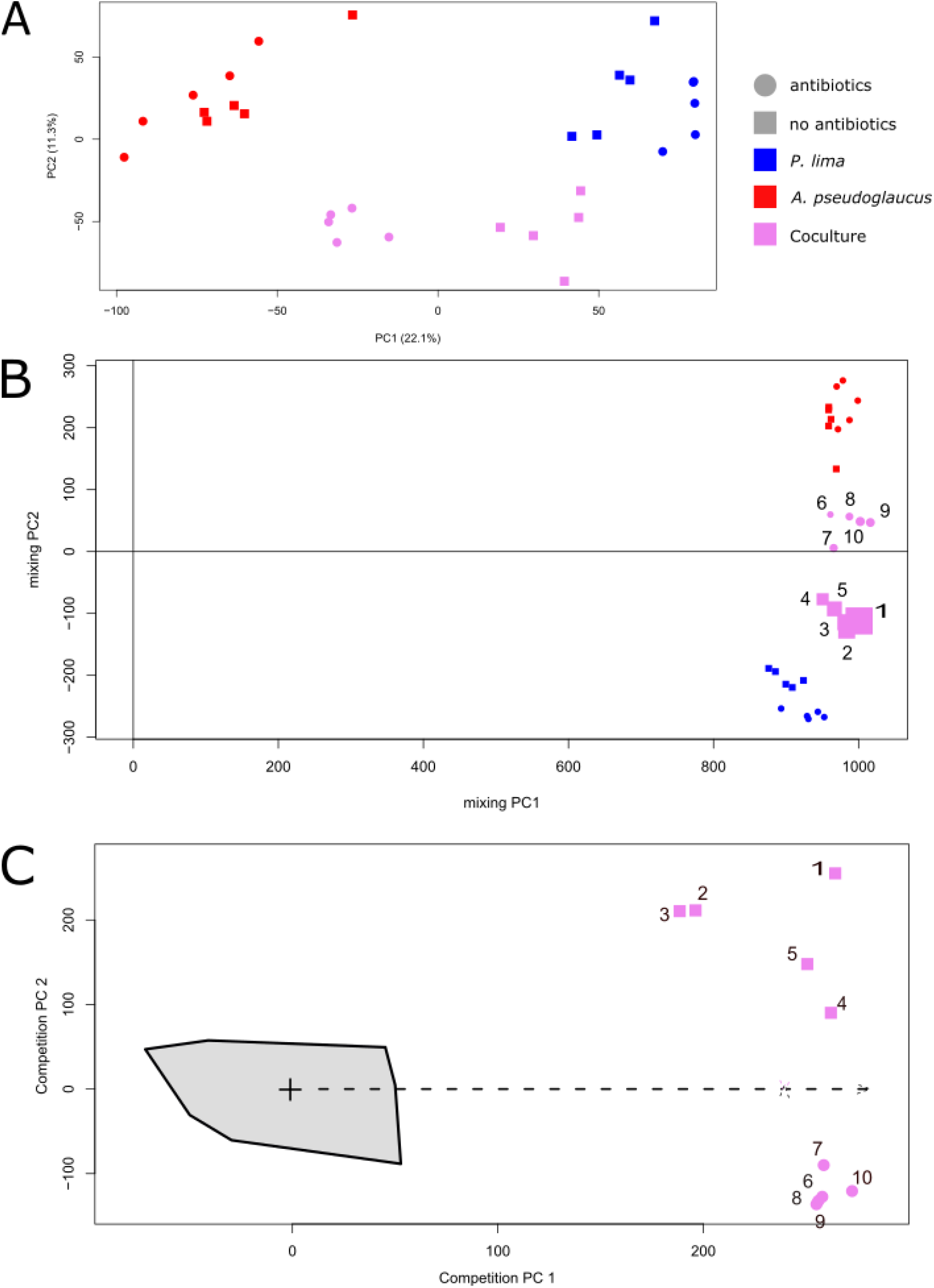
Unsupervised data analysis (n=5) of LC-HRMS data showing differences between *Prorocentrum lima, Aspergillus pseudoglaucus* and their co-culture samples, in absence or presence of antibiotics. A) Principal Component Analysis (log transformed data, pareto scaling); B) Mixing model using Projected Orthogonalised Encounter Monitoring (POChEMon) data analysis [50]. In the mixing model, the size of the co-culture samples reflects the amount of their unexplained variance within the mixing model (the larger the point is, the higher the unexplained variance is); C) Competition model using Projected Orthogonalised Encounter Monitoring (POCheMon) data analysis. The green zone corresponds to the projected co-culture domain within the competition model.

The PCA scores plot (Figure 2A) also highlighted a clustering of samples according to xenic and axenic status for each culture conditions. As signals related to antibiotics were removed from the data, the antibiotic clustering effect could mainly be linked to metabolome alterations due to the presence or absence of bacteria in the culture medium. In the presence of antibiotics, the chemical profiles of the *P. lima* monoculture corresponded only to compounds produced by the dinoflagellates, while without antibiotics, the chemical profiles reflect the metabolome of the microalgae with its associated bacteria. Greater differences between *P. lima* metabolomes under xenic and axenic conditions were observed in comparison to fungal ones under both conditions (Figure 2A), suggesting that the presence of antibiotics should alter more importantly the microalgal metabolite profiles than the fungal ones. Furthermore, an even greater impact of the xenic condition was observed within co-cultures. Also, the absence of antibiotics shifted co-cultured replicates toward the *P. lima* culture samples. This reflects the impact of *A. pseudoglaucus* on specialized metabolites produced by *P. lima* and its bacteria and the impact of microalgae (and associated bacteria) on *A. pseudoglaucus* metabolome.

Such results highlight the importance of bacteria in the chemical inertia of *P. lima* when confronted to *A. pseudoglaucus*. However, this unsupervised analysis did not point out the specific metabolome alteration induced by fungal presence as it encompasses every variation (microbiome or antibiotics derived variations). Thus, a more co-culture centric data mining strategy was used [31], called Projected Orthogonalised CHemical Encounter MONitoring (POChEMon) [49]. Such an approach uses an unsupervised mixing model, obtained by statistically mixing data from chemical profiles corresponding to strains grown alone. This yielded an *in-silico* co-culture model in absence of co-culture-related biotic stress. Then, projection of co-culture samples within this mixing model allowed, as for the PCA, to compare chemical profiles from the co-culture to monocultures. This approach further confirmed results from the PCA, showing that absence of antibiotics makes co-culture chemical profiles resemble more *P. lima* ones (Figure 2B). Similarly, the presence of antibiotics makes the chemical profiles of the coculture resemble more to the fungal profiles. Additionally, the POChEMon approach provided a global comparison between all co-culture experiments using the residual information, not represented in this mixing model (Figure 2B). The size of the co-culture samples when represented is directly related to unrepresented variance in the mixing model [49]. Here, this provided a strategy of unsupervised comparison between the two co-culture conditions (with or without antibiotics). Clearly, the chemical induction is stronger in presence of bacteria (larger residue) (Figure 2B). This further highlighted the importance of the microbiome in the co-culture specific chemical interaction, with more induction related to bacterial presence.

The co-culture induction pattern can be explored by interpreting the POChEMon competition model (Figure 2C) [49]. This representation corresponded to the unsupervised analysis through PCA of the co-culture sample residue when co-culture data were projected in the mixing model. Thus, the POChEMon approach allowed to identify co-culture related metabolome modifications in every sample. In Figure 2C, two co-culture patterns were observed and well separated corresponding to culture in presence or absence of antibiotics. This further confirmed the role of bacterial metabolites in co-culture, and also highlighted fungal-interaction specific compounds (residue in antibiotics treated samples). Interestingly the overall variability among co-culture replicates is lower in presence of antibiotics in comparison to their absence.

### Identification of co-culture up-regulated compounds highlights *Prorocentrum lima* toxin induction

As co-culture induction is a complex mechanism, different complementary supervised statistical approaches were used to highlight induced features [55]. Data were analysed univariately using Student test and multivariately using both orthogonal projection to latent structure discriminant analysis (OPLS-DA) and the POChEMon approach [49]. In all cases the top 20 most relevant features were selected with their retention time, *m/z* value and the statistical approach used. Finally, a set of 51 features were selected as over-produced (or *de novo* induced) during co-culture (Table 1 and Figure S3). Among those features, 14 were detected in the *P. lima* extracts, 6 in the *A. pseudoglocus* extracts and 34 were not detected in monocultures (ND). In addition, 12 and 42 features were observed in agar and liquid media, respectively. Most of the features were found in antibiotic-free samples as the mixing model emphasized (Figure 2B), but some of them were specifically found when antibiotics were added, hence confirming a fungal-algal-interaction related chemical induction.

**Table 1.**
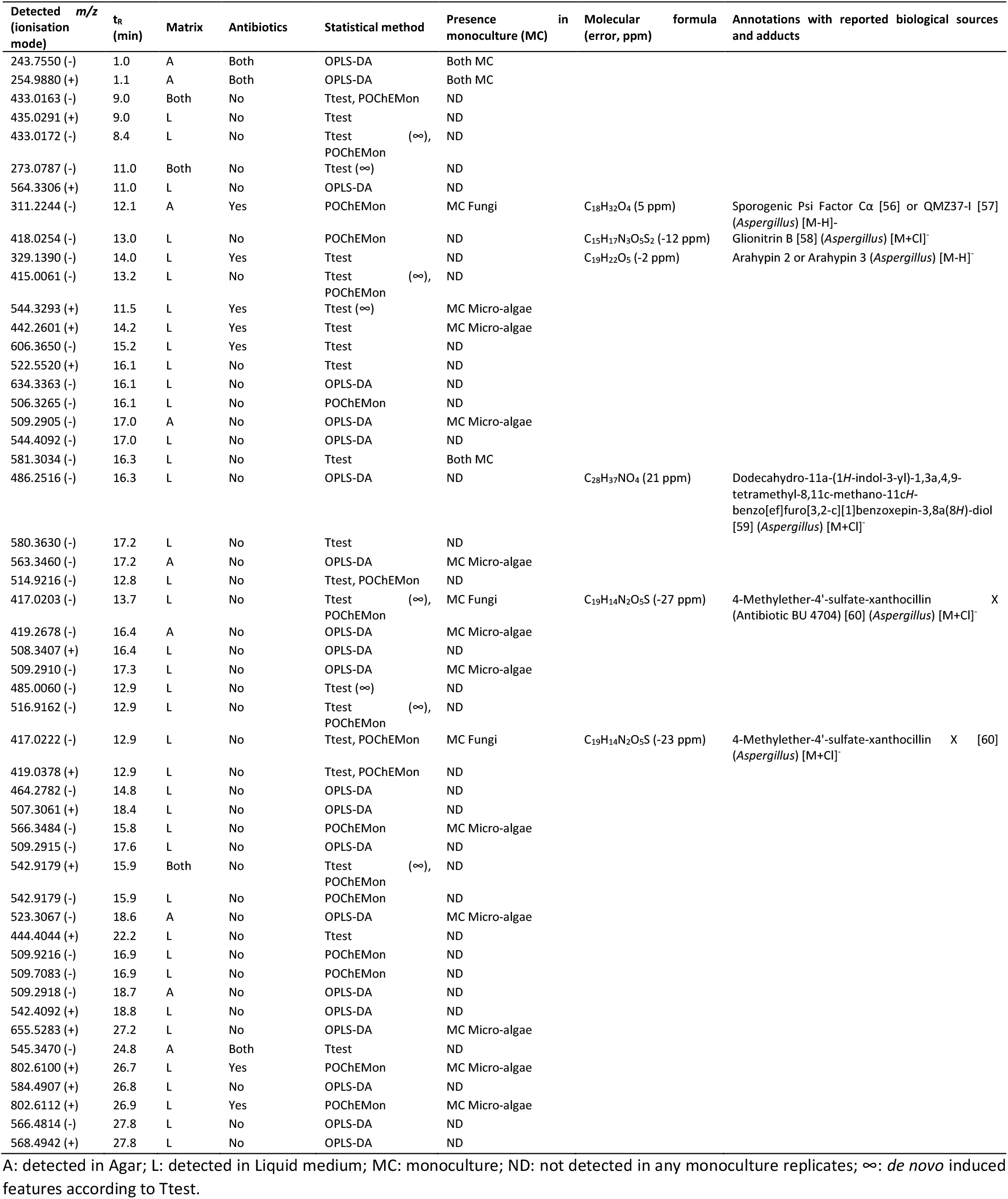
Co-culture associated features using univariate and multivariate data analyses using PocheRmon package. Features highlighted using Ttest method correspond to top 20 ones with fold changes higher than 2 with Student-test *p*-value below 5%. The OPLS-DA method select up-regulated features with VIP (variable importance in the projection) value higher than 2. Finally, the POChEMon method highlights features with SSrankProduct higher than 0.995. The drastic variable selection yielded low number of features highlighted by different analysis.

Based on high mass and spectral accuracy, the highlighted features were putatively annotated (Table 1 along with annotation accuracy based on Metabolomic standard initiative – MSI [61]). Considering the most significant features, none of the highlighted features was successfully annotated as *Prorocentrum* spp. metabolites. Furthermore, fungal metabolites annotation was only possible by extending the search to the clade *Aspergillus* section *restrictii*. As expected, only a very low number of features were identified.

Besides the previously highlighted features (Table 1), the two known toxins OA and DTX-1 usually produced by *P. lima* PL4V were also studied. Both toxins were found induced in co-culture with the Student test results in the LOP, with or without added antibiotics: OA (main ion detected at *m/z =* 827.458 [M+Na]^+^, t_R_ = 14.6 min) and DTX-1 (main ion detected *m/z =* 841.473 [M+Na]^+^, t_R_ = 17.1 min) with a fold change of 29.7 (*p*-value < 0.05) and 5 (*p*-value < 0.05), compared to monoculture conditions, respectively (they were ranked lower than 20). Okadaic acid was further confirmed using standard. No significant variations were observed in the solid phase profiles.

### Unprecedented physical contact was observed between *Aspergillus pseudoglaucus* and *Prorocentrum lima*

Visual, binocular and microscopic inspection of the co-culture samples revealed aggregation of *P. lima* within and around *A. pseudoglaucus* colonies (Figure 3AB). Only a limited number of free *P. lima* cells in biofilm stuck directly to the agar layer while most of biofilm is found directly in contact with *A. pseudoglaucus*. (Figure 3A). To further explore interaction morphology between *P. lima* PL4V and *A. pseudoglaucus* MMS1589, co-culture samples were explored using microscope. Surprisingly, an unusual physical interaction was observed between those organisms. Figure 3CD shows that the mycelium of *A. pseudoglaucus* seems to connect to the dinoflagellates at the flagella location. The red coloration of the connecting organelle indicated its fungal origin as Congo Red dye is known to highlight in red fungal membranes.

**Figure 3.**
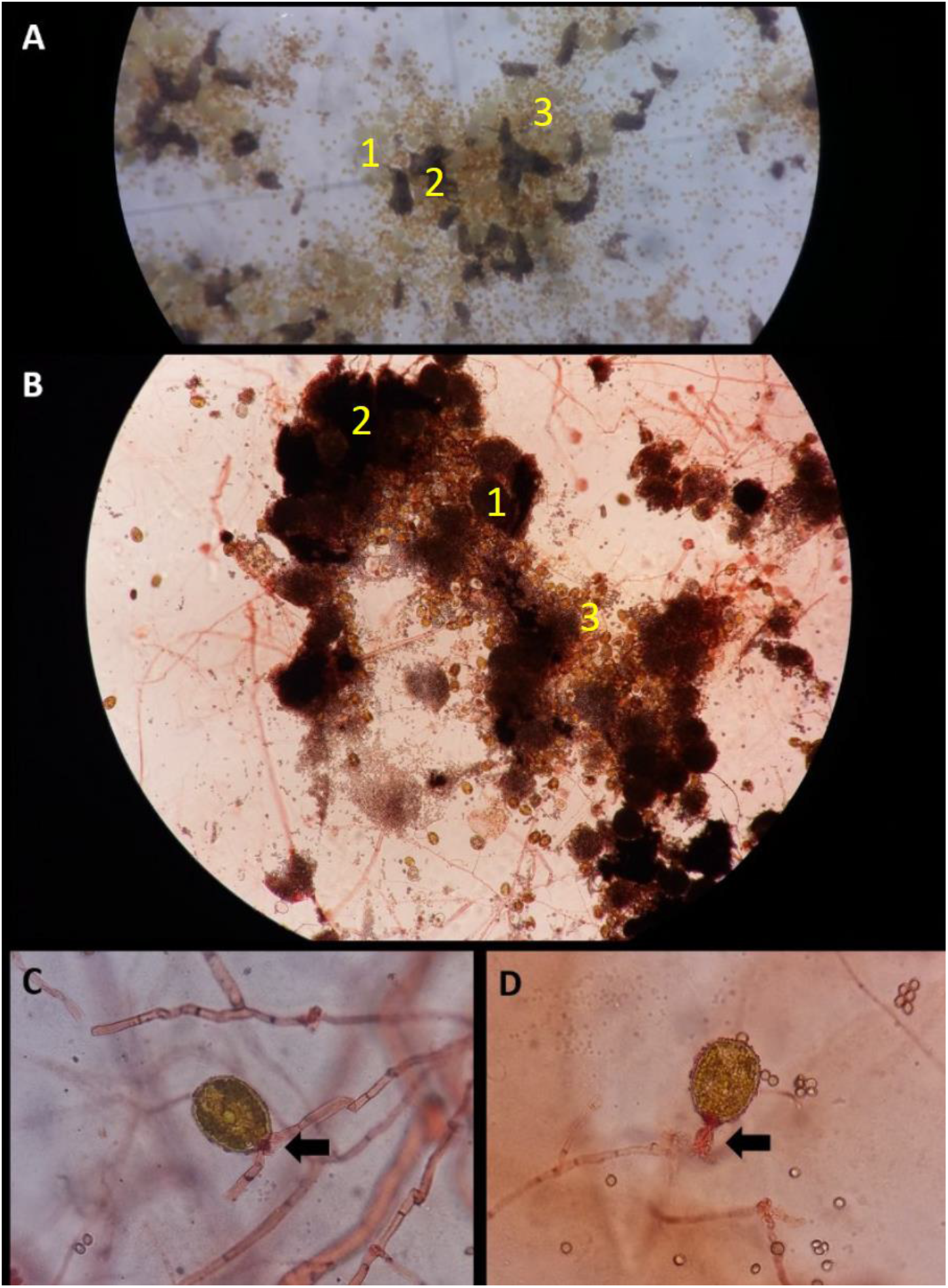
Observation of Fungal-microalgal interaction. AB) binocular observation of co-culture with aggregated cleistotheca (pale and round in A, brown in B) and asexual sporogenic spores (dark not round). Microalgae are oval and surround fungal elements. CDE) microscopic observation of co-culture with Congo Red dye (x400). The fungal mycelium is indicated (arrows) at proximity of the apex of the microalgae where the physical interactions occur.

## Discussion

Fungal role in the marine environment [62] is largely underexplored [63, 64]. However, fungal presence has been repeatedly observed, sometimes with rather large fungal biomasses [65–67]. The present study focuses on the interactions between the toxic dinoflagellate *P. lima* and an associated fungal strain. In fact, various fungal strains from the *Penicillium* and *Aspergillus* genera were isolated from the *P. lima* PL4V strain. The *A. pseudoglaucus* MMS1589 strain was selected to further explore chemical mediation between microalgae and fungi. Thus, a simple marine microcosm, compatible with fungal and benthic dinoflagellate life style, was devised to study such interaction. Specialized metabolites production during co-culture was studied by LC-HRMS metabolomics.

The differences of chemical alteration between axenic or xenic *P. lima* conditions clearly emphasize the effect of associated bacteria in the fungal/microalgal interaction (Figure 2). Interactions between *P. lima* and bacteria were shown to alter algal metabolites [68], and it is thus expected that in the present study the bacterial presence also significantly impacted the metabolite profile. Finally, this showed that, in addition to dinoflagellate signals, the microbiome metabolism is also altered. Up to now, chemical induction was mostly reported in fungal-fungal, bacterial-bacterial and fungal-bacterial co-cultures [34].

The chemical signal specifically related to *P. lima* and *A. pseudoglaucus* were tentatively identified. Some were reported to possess antimicrobial (asperchalasin E), antiviral (didemethylasteriquinone D, xanthocillin X) and anti-inflammatory activities (aspochalasin U). Furthermore, 5S,8R-Dihydroxy-9*Z*,12*Z*-octadecadienoic acid (*m/z* = 311.2244 at t_R_ = 12.11 min) was previously reported as a sporogenic factor of *Aspergillus* spp., which could reflect a fungal biotic stress adaptation behaviour. Unfortunately, among those highlighted features only a very limited number of compounds was putatively annotated. The reason may be that (1) frequently, microbial co-culture induces previously unreported compounds [34, 42] and that (2) metabolomes of toxic dinoflagellates are still largely unexplored (except toxins) [69]. Thus, a large effort in studying the metabolome of such organisms will be of great value to better understand their metabolism. Still, two induced compounds were unequivocally identified, OA and DTX-1, produced by *P. lima* PL4V. This is particularly interesting, as these toxins are largely related to *P. lima* toxicity by seafood contamination [70, 71]. In this experiment, signals from *A. pseudoglaucus* MMS1589 induces the over-production of such toxins. Similar results were reported with the domoic acid producer *Pseudo-nitzchia* where variation in bacterial community alters toxin levels [72, 73]. So far, there is very few evidence of toxin production alteration by associated microbiome [25, 74]. However, in this context, fungi should also be considered to play a significant role in such production regulation. Interestingly, OA and its derivatives (DTX-1 and 7-deoxyokadaic acid) are known to inhibit *Aspergillus niger* growth in a paper-disk assay [75]. Thus, OA and DTX-1 over-production may reflect that *P. lima* is tentatively controlling fungal growth; even though no evidence of fungal toxicity was observed in the presented study. Such inhibition was already observed during *P. lima* interaction with other microalgae [76]. Thus, *P. lima* toxin seems to be part of its global defence mechanism.

During the *P. lima* and *A. pseudoglaucus* interaction, aggregation was also observed (Figure 3AB). Similar aggregation followed by flocculation was previously related in biotechnology-focused fungal microalgal co-culture [77]. Such aggregation reflects the epiphytic nature of *P. lima* which needs to attach to objects in order to form biofilm [70]. This may also suggest nutrient exchange between both organisms which could be ease by their very close proximity. However, this needs more in-depth study to be proven in the present interaction between *A. pseudoglaucus* MMS1589 and *P. lima* PL4V. Such idea was recently explored in the synthetic association between the microalgae *Chlamydomonas reinhardtii* and the yeast *Saccharomyces cerevisiae* [78]. In this specific association, the two species were forced, under specific conditions, to mutually support each other’s growth. In a very recent study, during the tripartite interaction between *C. reinhardtii, Aspergillus nidulans* and *Streptomyces iranensis*, the fungal species was demonstrated to physically defend the microalgae from bacterial algicide compound azalomycin F by forming a protective lipid coat as observed by microscopy [79].

Besides aggregation and very unexpectedly, the microscopic observation of the interaction between *P. lima* and *A. pseudoglaucus* showed a very unusual physical contact where the fungal mycelium seems to be connected to the dinoflagellate at the flagella location (Figure 3). Such synthetic structured community formation is rather unusual in the literature and one may consider such contact as a very early stage of lichen formation [80, 81]. Lichens are classified as a microalgal-fungal symbiosis harbouring various other species, to form complex ecosystems [82]. *Aspergillus pseudoglaucus* was not previously mentioned as a lichen forming fungus [83]; however, some *Aspergillus* sp. were reported as endolichenic species having antimicrobial properties [84–86].

To conclude, this study raised many questions about microalgal/fungal interactions in the marine environment, especially in the context of toxic microalgae. Further studies are still needed to identify many of the metabolites induced either by *A. pseudoglaucus* MMS1589 or *P. lima* PL4V. It also would be interesting to further explore the mechanisms of such interactions and thus understand the physical contact observed and the OA induction reported here.

## Acknowledgements

The authors would like to acknowledge the technical assistance of Véronique Séchet and Florent Malo for maintaining and providing *P. lima* cultures. The authors thank the ThalassOMICS Metabolomics Facility – Plateforme Corsaire, Biogenouest, Nantes – for LC-HRMS analysis and metabolomics expertise. The authors would also like to acknowledge funding received from Ifremer and the regional research federation IUML for funding received for the Master studies of Alizé Bagot and Maud Chaigne.

