## Supplemental figures for "Deciphering microbiome impacts on fungal-microalgal interaction in a marine environment using metabolomics"

The authors declare no conflict of interest.

**Table S1.** Fungal strains isolated from *Prorocentrum lima* PL4V.

| Isolated fungal strain | Medium for isolation | Time needed to growth |
|---|---|---|
| MMS1589 Aspergillus pseudoglaucus | Dextrose Casein | 21 days |
| MMS1591 Aspergillus sp. | Dextrose Casein | 29 days |
| MMS1593 Penicillium sp. | Malt Extract | 27 days |
| MMS1594 Aspergillus sp. | Dextrose Casein | 27 days |
| MMS1596 Penicillium sp. | Malt Extract | 34 days |

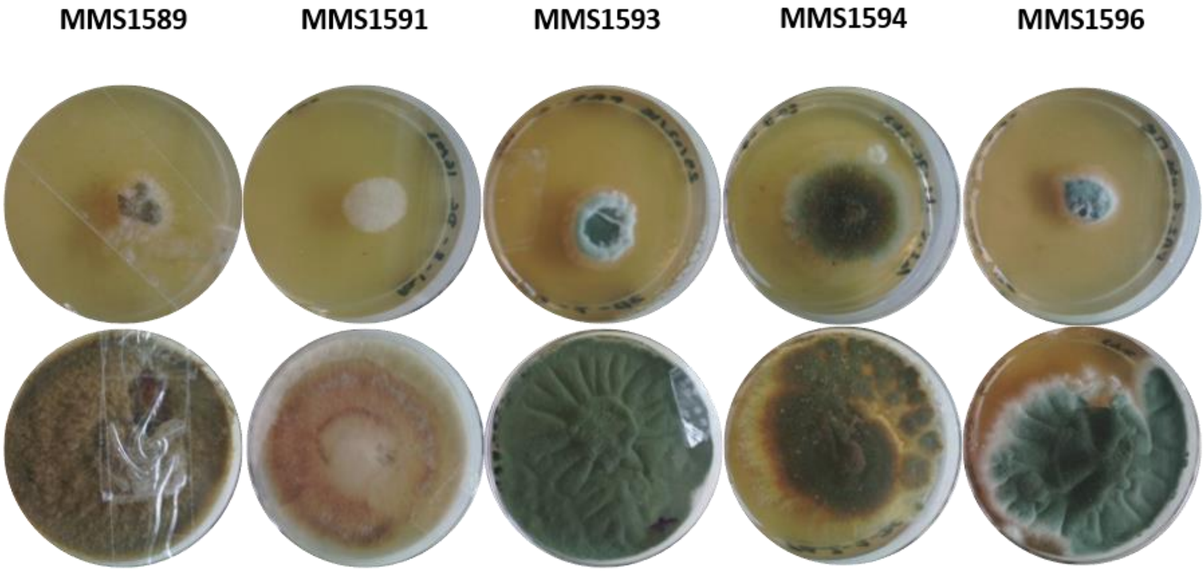


**Figure S1.** Morphology of fungal strains isolated from *Prorocentrum lima* PL4V

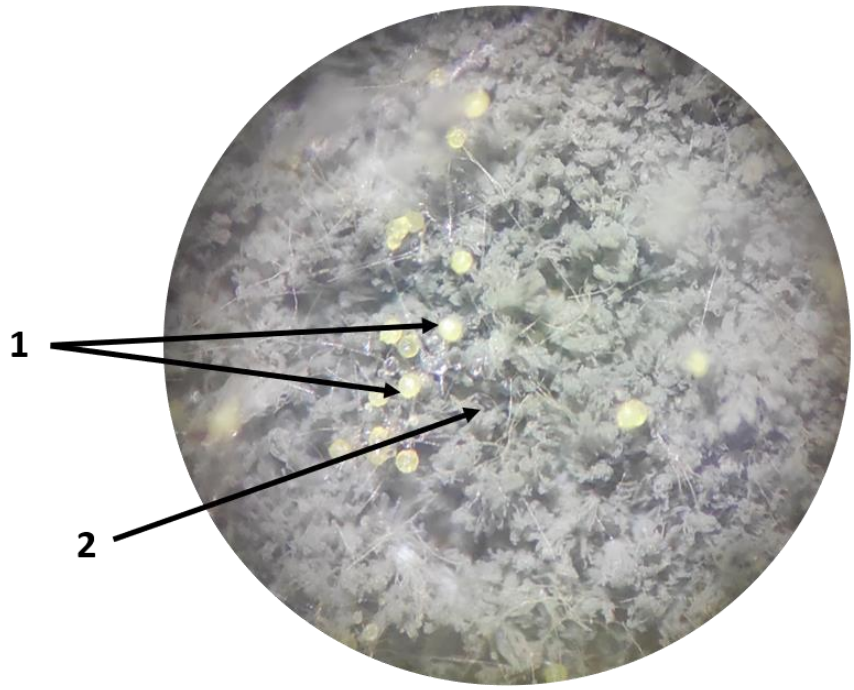


**Figure S2.** Morphology *of Aspergillus pseudoglaucus* MMS1589 (binocular view). Cleistotheca (1) and classic mycelium (2)

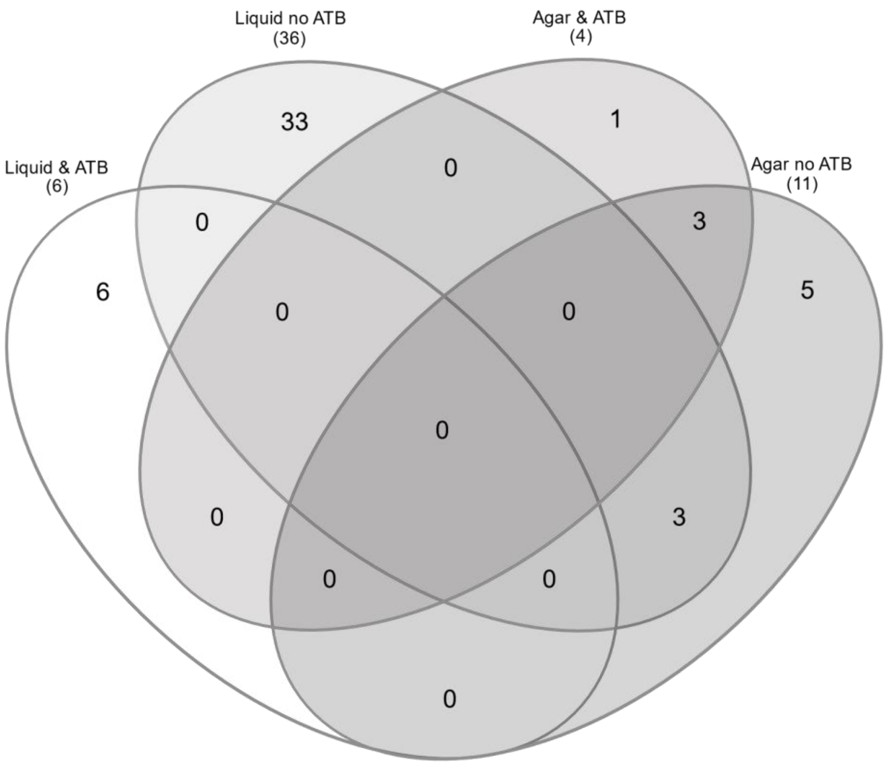


**Figure S3.** Venn Diagram [1] of feature distribution following matrix (liquid or solid) and antibiotic exposure.
